# Diversity and Distribution of Arrow Worms, Zooplankton Along the Mangaluru and Udupi Coast, Karnataka

**DOI:** 10.1101/2022.11.10.515975

**Authors:** Narshivudu Daggula, A.T. Ramachandra Naik, M. T Lakshmipathi, S. R Somashekara, N Manjappa, T Suresh

## Abstract

The present study was done to understand the abundance and species diversity of arrow worms, zooplankton in relation with hydrographical parameters and the spatial and temporal variations for a period of 16 months from October 2020 to March 2022. Monthly sampling was carried out at selected stations of coastal waters with average depth of 8m. The qualitative analyses revealed the presence of seventeen different groups of zooplankton. copepods, cladocerans are dominant group in zooplankton followed by chaetognaths, during the study period seven species of chaetognaths recorded viz, *Sagitta enflata, Sagitta robusta, Sagitta bedotii, Sagitta bipunctata, Sagitta ferox, Sagitta planktonis*, and *Sagitta elegans*. Among them, *Sagitta enflata* was the dominant species followed by *S*.*bedotii* and *S. robusta*. The richness, evenness and diversity of chaetognaths varied in the different stations and the maximum and minimum values also varied between stations, but in the most of the stations the minimum values were observed in the post-monsoon season and the highest values ranged between January to March. On comparison of the present abundance of chaetognaths with previous study, it was observed that the density of chaetognaths decreased compared to previous studies.

## INTRODUCTION

Chaetognatha or chaetognaths, meaning *bristle-jaws* are a phylum of predatory marine worms that are a major component of plankton worldwide, commonly known as arrow worms, about 20% of the known chaetognaths species are benthic, and can attach to algae and rocks mostly chaetognaths are transparent and torpedo shaped, but some deep-sea species are orange. They range in size from 2 to 120 millimeters (0.1 to 4.7 in). Among the zooplankton community, the chaetognaths are a relatively isolated phylum that includes a total of 209 species recorded in the world’s oceans, of which 29 have been recorded in the South Atlantic (Souza, C.S., Luz, J.A. and Mafalda Junior, P.O., 2014, Melo, D.C., *et al*., 2020). Chaetognath species are distributed worldwide, including the Pacific, Atlantic, Indian and Antarctic Oceans. They are found in most of the vertical realms spanning from the surface to the bottom of the ocean (Müller *et al*., 2019; WoRMS, 2022). At present, phylum Chaetognatha consists of over 150 defined and described species from all the marine ecosystems and depths of the world oceans (Giribet, G. and Edgecombe, G.D., 2020). Out of there 35 species belonging to 15 genera and 5 families, are recorded from the oceanic and deep-sea waters of India (Chandra, K. and Raghunathan, C., 2022). Chaetognaths are marine invertebrate predators that form a chief component of mesozooplankton community throughout the world oceans and are found in all marine ecosystems, including estuarine, coastal and oceanic waters, due to their body shape and quick darting locomotion in water. They are commonly called as “*arrow worms”* because of their darting movement (Srichandan, S., *et al*., 2015). The biomass of chaetognaths is estimated to be 10 to 30% of that of copepods in the world’s oceans (Sanvicente A. L., *et al*., 2020). Chaetognaths have two characteristics that make them particularly interesting from the oceanographic point of view. Firstly, they have proved to be good indicators of water masses and suitable for the study of the effects of Physico-chemical processes acting at the mesoscale on the dynamics and variability of zooplankton populations secondly, their role of predation as a decisive factor determining the structure of the marine planktonic food webs. They have a notable impact on zooplankton communities as one of the main predators of copepods, previously the works has been covered by (Pillai *et al*., 2000, Narale *et al*., 2015, Gupta *et al*., 2016. Srichandan, S., *et al*., 2015 and Karati, K.K., *et al*., 2018) in eastern and southwest coast of Arabian Sea.

## MATERIAL AND METHOD

The zooplankton sampling was carried out on board by vertical hauls performed at 10m depth. Zooplankton present in the columnar waters were collected using a Bongo net. The mouth area of the net is 0.25 m^2^ with a 250 cm long filtering cone. The mesh size of the filtering cone is 180 μm. For the collection at 10m depth, the net was hauled vertically leaving 2 meters safe depth. Zooplankton samples collected were transferred to a clean polythene bottle and preserved in a 5% formalin solution. Zooplankton samples were filtered through a 60 μm nylon bolting silk cloth. The samples were concentrated using a separating funnel and then re-suspended to make up the volume of up to 250 ml. For qualitative analysis, all chaetognaths were isolated from the total zooplankton samples. This was done by pouring the samples into a large petri dish. A table lamp was used for ease in observing the specimen. A 10x handheld Microscope was used to observe and separate the chaetognaths. The taxonomic observations were made on the preserved samples under a compound microscope (OLYMPUS CK X 41) microscope using the keys of previous investigations (Kapp, H. and Pierrot-Bults, A.C., 1991, Volkova, E. and Kudryavtsev, A., 2021.) For quantitative analysis every specimen present in the collected samples were counted. The species diversity and species evenness of phytoplankton and zooplankton were calculated by using the Shannon-Weiner diversity index (H1). Species richness and evenness (J1) were calculated using Pielou (Iqbal, M. M., *et. al* 2014) method. The analysis was performed using (Plymouth Routines in Multivariate Ecological Research) PRIMER-7 analytical package.

**Plate 1.**
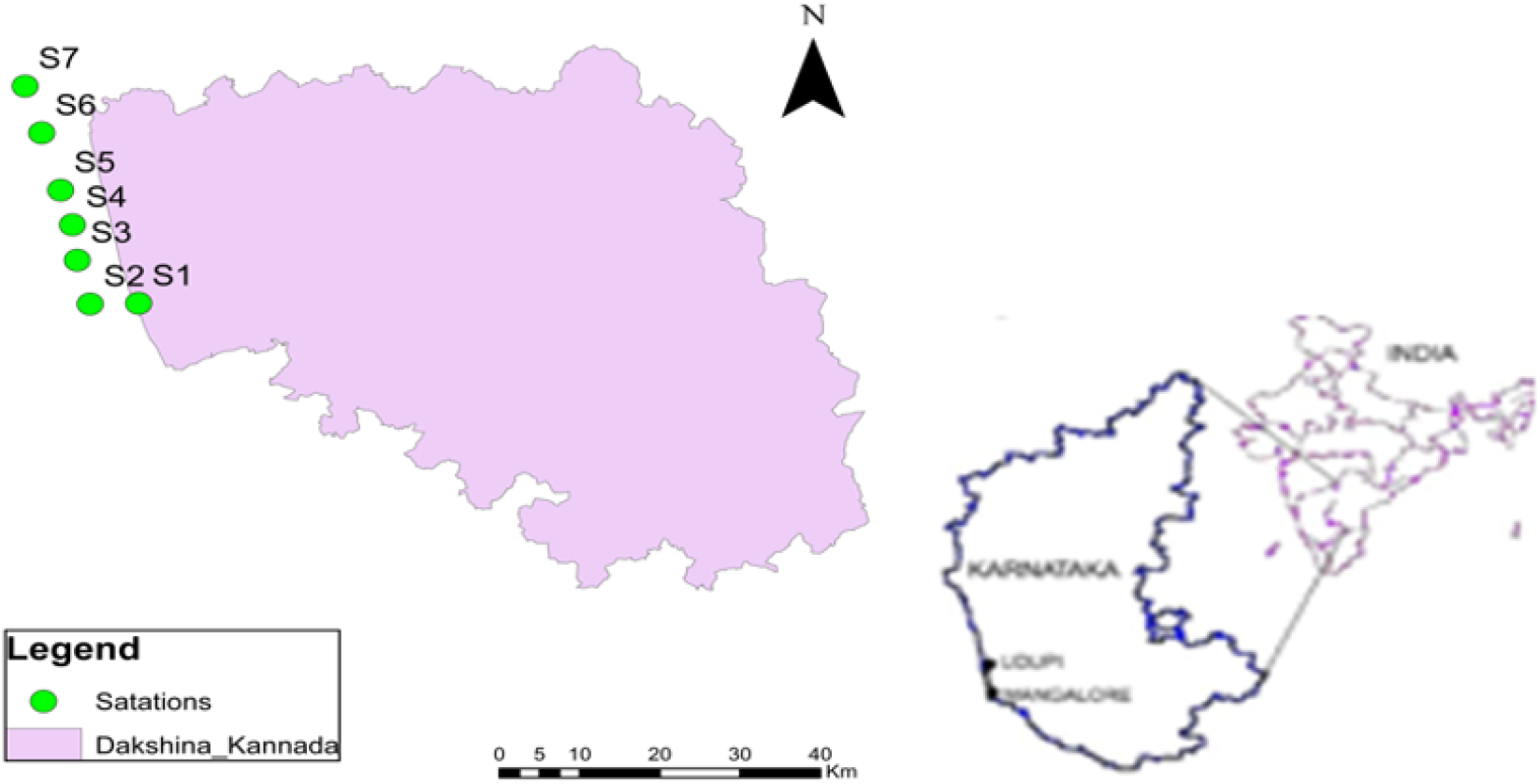
Map showing the location of sampling stations

## RESULTS AND DISCUSSION

In the present study, the species observed were *Sagitta enflata, S. robusta, Sagitta bedotii, S. bipunctata, S. ferox, S. planktonis* and *S. elegans*. The species composition was similar in most of the stations during the sampling period, but the species dominance varied with stations. Station Netravathi-Gurupura the total number of chaetognaths ranged from 20 to 365 individuals/m^3^ during the study period. The maximum (365 individuals/m3) number was observed during January, 2021 and the minimum (22 individuals/m3) was in October, 2021. The chaetognaths consisted of 5 species belonging to 1 genus. Among them, *Sagitta enflata* (871 individuals/m^3^) was the dominant species followed by *Sagitta robusta* (276 individuals/m^3^), and *Sagitta ferox* (141 individuals/m^3^). The *S. enflata* was recorded in all the samples and found most abundant of all the species observed. The second most dominant species found was *S. robusta* followed *S. bedotii*. Their abundance varied with different stations. *S. elegans* was observed only in the Netravathi-Gurupur waters and absent in other stations. *S. ferox* was found in all the stations except in coastal waters off Chitrapur. *S. enflata* is an epipelagic and cosmopolitan species found in many regions around the world (Iyyapparajanarasimapallavan *et al*., 2013). Station Panambur total number of chaetognaths ranged from 22 to 731 individuals/m^3^. The maximum (731 individuals/m^3^) was observed during May and the minimum (22 individuals/m^3^) was in October. The chaetognaths consisted of 6 species belonging to the same genera. Among them, *Sagitta enflata* (1850 individuals/m^3^), was the dominant species followed by *Sagitta robusta* (372 individuals/m^3^) and *Sagitta bedotii* (216 individuals/m^3^). Station Panambur the total number of chaetognaths ranged from 28 to 382 individuals/m^3^. The maximum (382 individuals/m^3^) was observed during December and the minimum (28 individuals/m^3^) was October. The chaetognaths consisted of 5 species belonging to same genera. Among them, *Sagitta enflata* (1262 individuals/m^3^) was the dominant species followed by *Sagitta bedotii* (184 individuals/m^3^) and *Sagitta robusta* (162 individuals/m^3^). Station Chitrapura the total number of chaetognaths were ranged from 22 to 802 individuals/m^3^. The maximum (802 individuals/m^3^) was observed during April and the minimum (22 individuals/m^3^) was October. The chaetognaths consisted of 5 species belonging to the single genera. Among them, *Sagitta enflata* (1441 individuals/m^3^) was the dominant species followed by *S. bedotii* (624 individuals/m^3^) and *S. robusta* (186 individuals/m^3^). It is a temperate and warm water species (Muthukumaravel, 2021). Hence, they are found in abundance in the present study period, according to B.K Sahu (2013) (Velmurgan, P., 2014), S. *enflata* was more abundant in coastal regions. *S. bedotii, S*.*ferox* and *S. robusta* are Indo-Pacific species that are epipelagic (Srichandan, S., *et al*., 2015). *S. bipunctata* is an oceanic, cosmopolitan temperate to tropical water. Chaetognath was found year-round. Seasonal and station-wise variation showed that the abundance of chaetognath varied with stations and months. In October, the abundance was very less and the sample mainly constituted of larval form. There was a gradual increase in the count from October to November reaching the peak in December. There was decrease in February with a rise again in March following a gradual decrease in May. Hence, there was bimodal oscillation in the season-wise variation of the abundance of the chaetognaths. The highest abundance was observed in the pre monsoon season (March to May). This could be attributed to biological factors like growth, reproduction and increased metabolic activity with an increase in temperature and salinity. The increase in the number of *S. bedotiid* and *S. robusta* which was lesser in the previous months contributed to the increase in the abundance in this period, the maximum chaetognaths were observed during May and the minimum was October. Among them, *Sagitta enflata* was the dominant species followed by *S. robusta*. The richness, evenness and diversity of chaetognath varied in the different stations and the maximum and minimum values also varied between stations. But in most of the stations, the minimum values were observed October and the highest value ranged in between January to March. Similarly, community structure and distribution of chaetognaths were investigated by Purushothaman, A., *et al*. (2021) along the upwelled and non-upwelled waters of the south-eastern Arabian sea (SEAS), Chaetognaths belonging to 10 genera were identified which genus *Flaccisagitta* (54%) made the most dominant group along the entire study area followed, in order of abundance, by *Serratosagitta* (20%), *Mesosagitta* (18.2%), *Sagitta* (12.3%), *Ferosagitta* (11%) and *Krohnitta* (6.4%). *Flaccisagitta* were observed to be abundant in the upwelled waters along with *Pterosagitta, Serratosagitta, Sagitta, Krohnitta* and *Ferosagitta* whereas genus *Mesosagitta* dominated the non-upwelled waters of northern transects

**Fig. 1.**
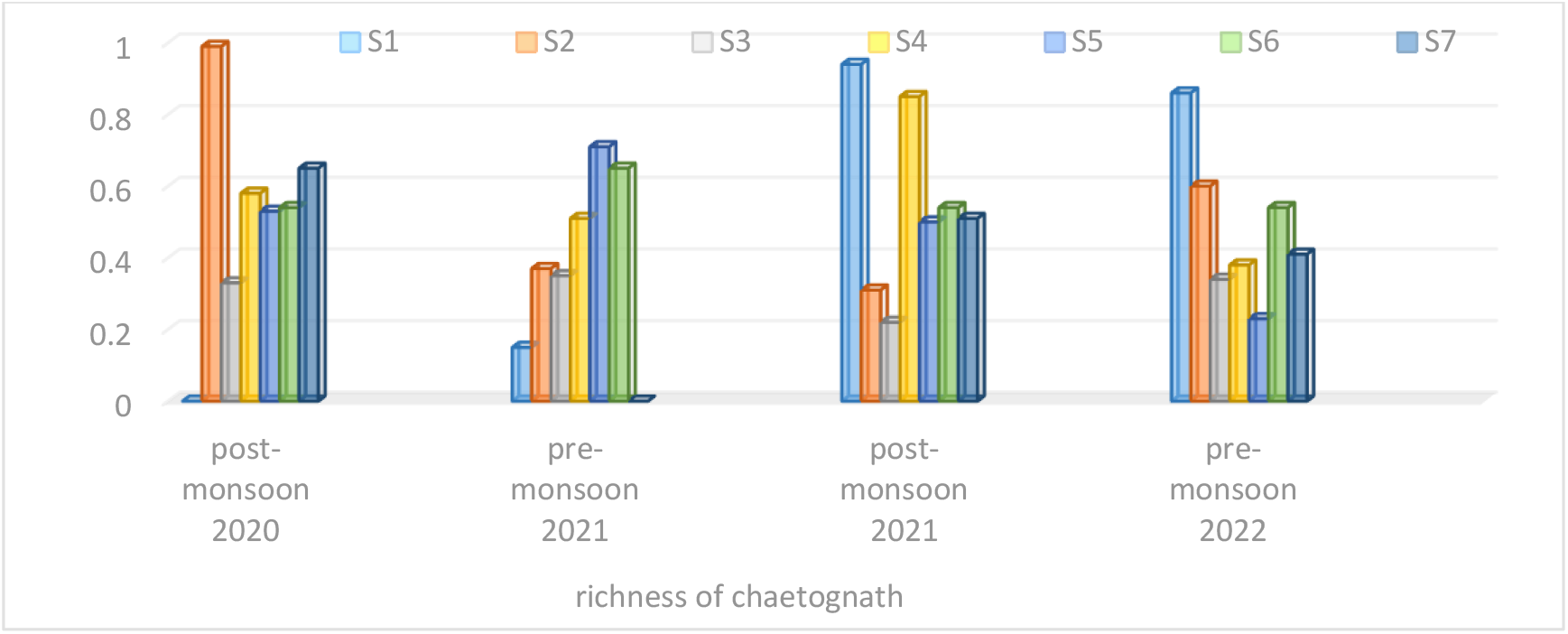
Seasonal variations in the richness of chaetognaths at selected stations.

**Fig. 2.**
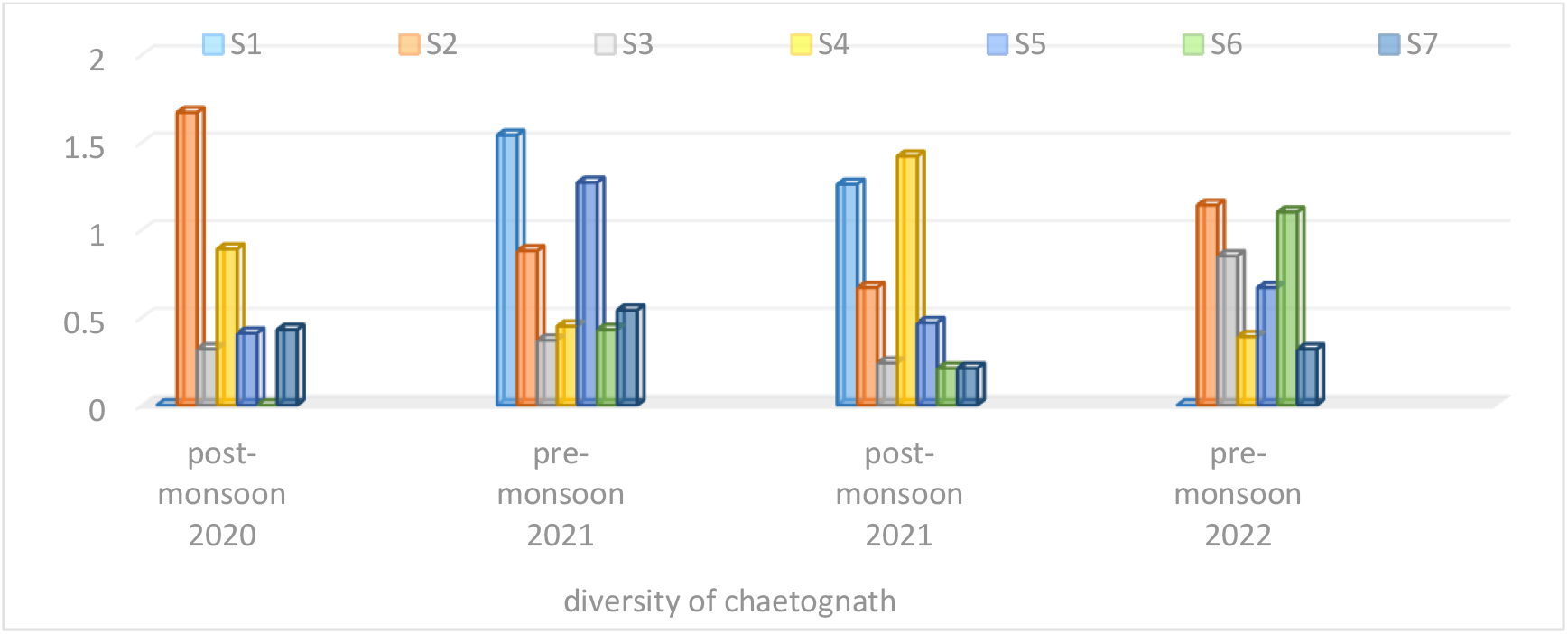
Seasonal variations in the diversity of chaetognaths at selected stations.

**Fig. 3.**
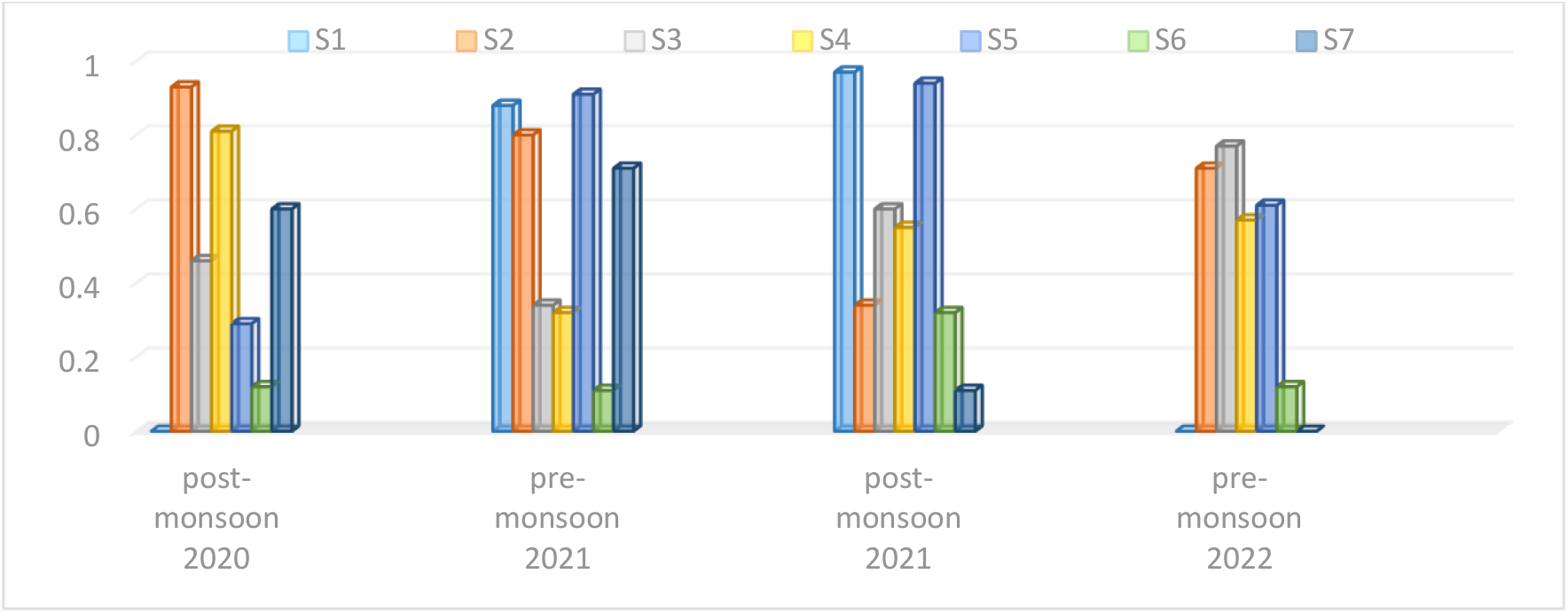
Seasonal variations in the evenness of chaetognaths at selected stations.

Besides the onset of the monsoon season also affected the abundance of the chaetognaths, also the increase in salinity seemed to favour the abundance of the chaetognaths. The Extinction coefficient was also higher during these months which indicated higher zooplankton in these months. Chaetognatha being highly carnivorous feeding, especially on copepods found food in high quantities during this period. The abundance in post-monsoon season during November to December can be attributed to the effect of upwelling which brings nutrient-rich water to the surface increasing the productivity of the plankton (Nair *et al*., 1991). Similar readings were observed by Gupta *et al*. (1988), Shwetha (2010), Shruthi (2015) and Karati, K.K., 2021, who observed higher abundance in post-monsoon and pre-monsoon season. The maximum (0.99) richness was observed November at station 2 (coastal waters off Thannirbhavi) and minimum (0) was observed in the month of October in all the stations except at station S4. due to high productivity, Chaetognatha are primarily marine and hence their relative higher abundances in the macro-tidal estuaries can be linked to their specific preferences for a high-saline environment (Júnior *et al*., 2019) although there are marked seasonal density fluctuations due monsoons in the west coast of India (Vineetha *et al*., 2015, Vishnu Radhanan *et al*., 2015, Gupta *et al*., 2016) The maximum (0.97) evenness was recorded in February at station S2 (coastal waters off Thannirbhavi) and the minimum (0) was observed in October in all the station except station S4. (Kusum *et al*., 2011, Nair, V.R., et al., 2015, Karati, *et al*., 2022) recorded lower evenness in November to February. Similarly (Lathasumathi, *et al*., 2017) studied the seasonal variation of community composition of zooplankton in the Palk strait, as the present study is a seasonal quantitative approach of chaetognaths dynamics, the causes of their Spatio-temporal variations can be the future line investigation. But what controls the season-wise and station wise fluctuation in Chaetognaths abundance and species composition should be further studied in relation to environmental and anthropogenic factors.

**Table.**
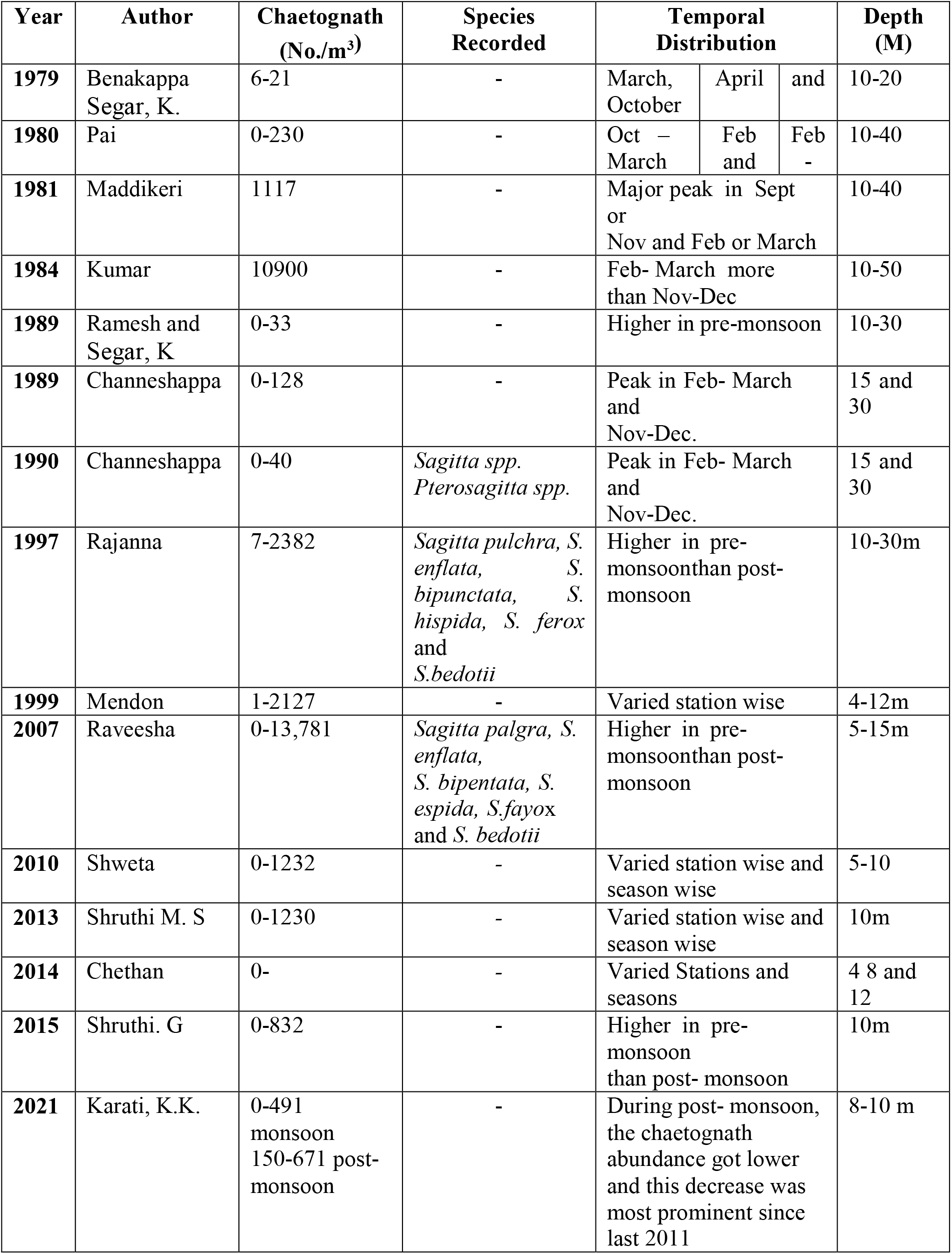
Monthly distribution of chaetognath (no./m3) from 1979 to 2022 at coastal waters off Mangaluru

## CONCLUSION

Chaetognaths maximum density was observed during the post-monsoon, among them, *Sagitta enflata* was the dominant species followed by *S. robusta* then *S. ferox*. Station Panambur, the maximum chaetognaths were observed during the summer and the minimum was in post-monsoon. Among them, *Sagitta enflata* was the dominant species followed by *S. robusta* and *S. bedotii*. Station Panambur, the maximum chaetognath was observed during summer and the minimum was in post-monsoon among them, *Sagitta enflata* was the dominant species followed by *S. bedotii* and *S. bipunctata*. Station chitrapura, the maximum chaetognaths was observed during post monsoon. Among them, *Sagitta enflata* was the dominant species followed by *S. bedotii* and *S. robusta*. Station Padubidri, the maximum chaetognath was observed during post-monsoon and the minimum was in the month of pre-monsoon. Among them, *Sagitta enflata* was the dominant species followed by *S*.*bedotii* and *S. robusta*. The richness, evenness and diversity of chaetognaths varied in the different stations and the maximum and minimum values also varied between stations but in most of the stations, the minimum values were observed in the post-monsoon season and the highest value ranged in January to March on comparison of the present abundance of chaetognaths with the previous study, it was observed that the density of chaetognaths decreased compared to previous studies.

## REFERENCES

Ajithkumar, T.T., Thangaradjou, T. and Kannan, L., 2006. Physico-chemical and biological properties of the Muthupettai mangrove in Tamil Nadu. J.Mar. Biol. Ass. India, 48: 131–138.

Baliarsingh, S.K., Srichandan, S., Naik, S., Sahu, K.C., Lotliker, A.A. and Baxter, E.J., Rodger, H.D., Mcallen, R. and Doyle, T.K., 2011. Gill disorders in marine farmed salmon: Investigating the role of hydrozoan jellyfish. Aquaculture Environment Interactions, 1: 245–57.

Chakrapany, S., 2012. Studies on Marine invertebrates-Scyphomedusae of the India and adjoining seas. Ph.D. thesis, Uni. Mad., Ind.

Chandra, K. and Raghunathan, C., 2022. Status, Issues, and Challenges of Biodiversity: Invertebrates. In Biodiversity in India: Status, Issues and Challenges (pp. 77–117). Springer, Singapore.

Damotharan, P.N., Perumal, VENGADESH., Arumugam, M., Perumal, P., Hamner, W.M. and Dawson, M.N., 2009. A review and synthesis on thesystematics and evolution of jellyfish blooms: advantageous aggregations and adaptive assemblages. Hydrobiologia, 616: 161–191.

DE LOS Rios, P., Kalaiarasi, M., Paul, P., Lathasumathi, C. and Stella, C., 2019. Monthly variations in crustacean zooplankton abundances in Sundarapandian Pattinam and Manamelkudi in the Palk Strait, India (9-10° N, Arabian Sea). Crustaceana, 92(3), pp.295–306.

Giribet, G. and Edgecombe, G.D., 2020. The invertebrate tree of life. Princeton University Press.

Gupta, G.V.M., V. Sudheesh, K.V. Sudharma, N. Saravanane, V. Dhanya, K.R. Dhanya, M. Sudhakar, and S.W.A. Naqvi., 2016. Evolution to decay of upwelling and associated biogeochemistry over the southeastern Arabian Sea shelf. Journal of Geophysical Research and Biogeosciences, 121, 159–175, 2016.

Iqbal, M.M., Islam, M.S. and Haider, M.N., 2014. Heterogeneity of zooplankton of the Rezukhal Estuary, Cox’s Bazar, Bangladesh with seasonal environmental effects. International Journal of Fisheries and Aquatic Studies, 2(2), pp.275–282.

Iyyapparajanarasimapallavan, G., Kumar, P.S., Kumar, C.P., Jalal, K.C.A., Kamaruzzaman, B.Y. and John, B.A., 2013. Distribution and abundance of gelatinous zooplankton along Tamil Nadu coastal waters. Journal of Biological Sciences, 13(1), pp.18–25.

Kapp, H. and Pierrot-Bults, A.C., 1991. The biology of chaetognaths (pp. 137–147). Oxford, UK: Oxford University Press.

Karati, K.K., Ashadevi, C.R., Harikrishnachari, N.V., Valliyodan, S., Kumaraswami, M., Naidu, S.A. and Ramanamurthy, M.V., 2021. Hydrodynamic variability and nutrient status structuring the mesozooplankton community of the estuaries along the west coast of India. Environmental Science and Pollution Research, 28(31), pp.42477–42495.

Karati, K.K., Vineetha, G., Devassy, R.P., Al-Aidaroos, A.M. and El-Sherbiny, M.M., 2022. Role of Ecohydrographical Barriers on the Spatio-Temporal Distribution of Chaetognath Community in the Gulf of Aqaba during Summer. Water, 14(5), p.822.

Kumar, T.S., 2013. Distribution of hydro-biological parameters in coastal waters off Rushikulya estuary, East coast of India: A pre-monsoon case study. Pak. J. Biol. Sci., 16(16): 779–787.

Lathasumathi, C., Escalante, P.D.R., Kalaiarasi, M. and Stella, C., 2017. Seasonal variation of community composition of zooplankton in the Palk strait,(9-10° N, Arabian sea, India). Bulletin de la Société Royale des Sciences de Liège.

Melo, D.C., Lira, S.M., Moreira, A.P.B., Freitas, L., Lima, C.A., Thompson, F., Bertrand, A., Silva, A.C. and Neumann-Leitao, S., 2020. Genetic diversity and connectivity of Flaccisagitta enflata (Chaetognatha: Sagittidae) in the tropical Atlantic ocean (northeastern Brazil). PloS one, 15(5), p.e0231574.

Müller, C.H, Harzsch, S. and Perez, Y., (2019) 7. Chaetognatha (Handbook of Zoology). Miscellaneous Invertebrates, De Gruyter, Germany, 282 pp. https://doi.org/10.1515/9783110489279-007

Muthukumaravel, K., Pradhoshini, K.P., Vasanthi, N., Raja, T., Jaleel, M.A., Arunachalam, K.D., Musthafa, M.S., Ayyamperumal, R., Karuppannan, S., Rajagopal, R. and Alfarhan, A., 2021. Assessment of seasonal variation in distribution and abundance of plankton and ichthyofaunal diversity in relation to environmental indices of Karankadu Mangrove, South East Coast of India. Marine Pollution Bulletin, 173, p.113142.

Nair, V.R., Kusum, K.K., Gireesh, R. and Nair, M., 2015. The distribution of the chaetognath population and its interaction with environmental characteristics in the Bay of Bengal and the Arabian Sea. Marine Biology Research, 11(3), pp.269–282.

Narale, D.D., P.D. Naidu, A.C. Anil, and S.P. Godad, 2015. Evolution of productivity and monsoonal dynamics in the eastern Arabian sea during the past 68 ka using dinoflagellate cyst records. Palaeogeography Palaeoclimatology and Palaeoecology, 435: 193–202.

Peterson, W.T., 2009. Copepod species richness as an indicator of long-term changes in the coastal ecosystem of the Northern California Current. Cal COFI Report, 50: 73–81.

Pillai, V.N., V.K. Pillai, C.P. Gopinathan, and A. Nandakumar, 2000. Seasonal variations in the physico-chemical and biological characteristics of the eastern Arabian Sea. Journal of Marine Biology Association of India, 42: 1–20.

Pratap, G. V., and Babu, K. R., 2015. Seasonal variations of physic-chemical characteristics of water samples in Sarada and Varaha estuarine complex, East coast of India. Europea Academic Research. 3 (1): 472–486.

Purushothaman, A., Thomas, L.C., Nandan, S.B. and Padmakumar, K.B., 2021. Influence of upwelling on the chaetognath community along the Southeastern Arabian Sea. Wetlands Ecology and Management, 29(5), pp.731–743.

Ravichelvan, R., Ramu, S. and Anandaraj, T., 2015. Seasonal variations of water quality parameters in southeast coastal waters of Tamil Nadu, India. Int. J. Modn. Res. Revs., 3(10): 826–829.

Rissik, D. and Suthers, I., 2008. The importance of plankton. In: Suthers, I.M., Rissik, D., (Eds.), Plankton: a guide to their ecology and monitoring for water quality. CSIRO Publishing, Colling wood. pp 1–14

Sahu, B.K., Baliarsingh, S.K., Srichandan, S. and Sahu, K.C., 2013. Seasonal variation of zooplankton abundance and composition in Gopalpur creek: a tropical tidal backwater, east coast of India. Journal of the Marine Biological Association of India, 55(1), pp.59–64.

Sanvicente-Anorve, L., Sierra-Zapata, S., Lemus-Santana, E., Ruiz-Boijseauneau, I. and Soto, L.A., 2020. Feeding of Flaccisagitta enflata (Chaetognatha) upon copepods in the southern Gulf of Mexico. Cah. Biol. Mar, p.61.

Satpathy, K.K., Mohanty, A.K and Natesan, U., 2010. Seasonal variation in physicochemical properties of coastal waters of Kalpakkam, east coast of India with special emphasis on nutrients. Environ. Monit. Assess., 164: 153–171.

SHRUTHI, 2015. Species diversity of diatoms in relation to hydrographical characteristics in the coastal waters off Dakshina Kannada and Udupi district. M.F.Sc., thesis, Univ. Agril. Sci., Bangl, India.

SHWETA, 2010. Meiofauna in relation to sediment characteristics in Mulki estuary, Dakshina Kannada. M.F.Sc., thesis. Kar. Vet. Ani. Fish. Sci. Univ., Bidar.

Souza, C.S., Luz, J.A. and Mafalda Junior, P.O., 2014. Relationship between spatial distribution of chaetognaths and hydrographic conditions around seamounts and islands of the tropical southwestern Atlantic. Anais da Academia Brasileira de Ciências, 86, pp.1151–1165.

Shruthi, M.S. and Rajashekhar, M., 2013. Ecological observations on the phytoplankton of Nethravati-Gurupura estuary, south west coast of India. Journal of the Marine Biological Association of India, 55(2), pp.41–7.

Srichandan, S., Sahu, B.K., Panda, R., Baliarsingh, S.K., Sahu, K.C. and Panigrahy, R.C., 2015. Zooplankton distribution in coastal water of the North-Western Bay of Bengal, off Rushikulya estuary, east coast of India.

Suthers, I.M. and Rissik, D., 2008. Plankton. A guide to their ecology and monitoring for water quality, pp.115–132.

Vega-Pérez, L.A. and Schinke, K.P., 2011. Checklist of chaetognaths phylum from São Paulo State, Brazil. Biota Neotrop, 11(1): 1–9.

Vega-Pérez, L.A. and Schinke, K.P., 2011. Checklist of chaetognaths phyllum from São Paulo State, Brazil. Biota Neotropica, 11, pp.541–550.

Velmurgan, P., Srinivasan, A., Athithan, A., Padmavathy, P., Manimegalai, D. and Anand, T., 2014. Diversity and seasonal variations of plankton in coastal waters receiving salt pan effluents off Thoothukudi, south-east coast of India. Indian Journal of Fisheries, 61(4), pp.60–67.

Vijayalakshmi, S. and Balasubramanian T., 2010. Studies on Zooplankton ecology from Kodiakkarai (Point Calimere) Coastal Waters (South East Coast of India). Res.J. Biol. Sci., 5(2): 187–198.

Vineetha, G., N.V. Madhu, K.K. Kusum, and P.M. Sooria, 2015. Seasonal dynamics of the copepod community in a tropical monsoonal estuary and the role of sex ration in their abundance pattern. Zoological Studies, 54: 54.

Vishnu Radhan, R., S. Jerome, S. Ebin, P. Vethamony, P. Shirodkar, Z. Zainudin, and S. Shirodkar, 2015. Southwest monsoon influences the water quality and waste assimilative capacity in the Mandovi estuary (Goa State, India). Chemical Ecology 31: 217–234.

Volkova, E. and Kudryavtsev, A., 2021. A morphological and molecular reinvestigation of Janickina pigmentifera (Grassi, 1881) Chatton 1953–an amoebozoan parasite of arrow-worms (Chaetognatha). International Journal of Systematic and Evolutionary Microbiology, 71(11), p.005094.

WoRMS., Editorial Board., 2022. World Register of Marine Species. https://www.marinespecies.org.

Júnior MN, Da Costa BS, Martinez TA, Brandini FP, Miyashita LK., 2019. Diversity of gelatinous zooplankton (Cnidaria, Ctenophora, Chaetognatha and Tunicata) from a subtropical estuarine system, southeast Brazil. Mar Biodivers 49:1283–1298.

Segar, K. and Hariharan, V., 1989. Seasonal distribution of nitrate, nitrite, ammonia and plankton in effluent discharge area off Mangalore, west coast of India.

